# Activation of the Spx redox sensor counters cysteine-driven Fe(II) depletion under disulfide stress

**DOI:** 10.64898/2025.12.03.692063

**Authors:** Abigail G. Hall, Abdulelah A. Alqarzaee, Aliyah J. Collins, Sasmita Panda, Svetlana Romanova, Sujata S. Chaudhari, Andrew J. Monteith, Dorte Frees, Vinai C. Thomas

## Abstract

In many low G+C Gram-positive bacteria, the global regulator Spx helps maintain thiol homeostasis during disulfide stress, when protein thiols form aberrant disulfide bonds that can lead to misfolding and oxidative damage. Spx-dependent gene expression is triggered when an intramolecular disulfide bond forms between two cysteines in its redox switch. Surprisingly, some Spx functions persist even in the absence of an active redox switch, highlighting the need to better understand the physiological significance of maintaining this regulatory feature. Here, we utilize a *spx*^C10A^ mutant that encodes a redox-insensitive Spx variant to study the role of the Spx redox switch in *Staphylococcus aureus*. We show that the *spx*^C10A^ mutant is hypersensitive to diamide-induced disulfide stress and exhibits widespread transcriptional dysregulation of genes that contribute to thiol maintenance and disulfide repair. Remarkably, the *spx*^C10A^ mutant rapidly adapts to disulfide stress by increasing its intracellular pool of L-cysteine (L-Cys) through enhanced uptake, which helps restore a reduced intracellular environment. However, during this process increased L-Cys inadvertently depletes cytosolic Fe(II), leading to growth inhibition of the *spx*^C10A^ mutant. Finally, we show that the Spx-dependent control of intracellular L-Cys is critical for *S. aureus* survival when it encounters human neutrophils. Overall, these findings suggest that staphylococcal adaptation to disulfide stress through intracellular L-Cys accumulation imposes significant fitness costs that *S. aureus* overcomes by rapid regulatory control of thiol homeostasis through a functional Spx redox switch.

**Significance:** All cells have a pool of low molecular weight thiols, such as cysteine, glutathione, bacillithiol, and coenzyme A, to maintain redox balance under oxidative and disulfide stress. Among these, cysteine is a very effective thiol but is highly reactive, and its intracellular concentration must be tightly regulated. In *S. aureus*, we found that cysteine accumulates intracellularly during disulfide stress and if left unchecked, can inadvertently deplete cytosolic Fe(II), leading to growth inhibition. To prevent cysteine toxicity, *S. aureus* activates the global regulator Spx, which rapidly induces genes that restore thiol homeostasis and limits cysteine accumulation.

## Introduction

Bacterial pathogens that infect human tissues must overcome multiple host defenses that restrict their growth (1, 2). Among these host defenses, small reactive molecules such as reactive oxygen species (ROS), reactive nitrogen species (RNS), reactive chlorine species (RCS), reactive sulfur species (RSS), and reactive electrophilic species (RES) inhibit bacterial growth by disrupting intracellular redox balance, usually driving cells toward a more oxidized state, and damaging macromolecules essential for normal physiological processes. (3). In addition, host-derived short-chain and polyunsaturated fatty acids can also perturb metabolism and promote the accumulation of ROS and RES, respectively, compounding oxidative stress (4–6). To survive in this hostile environment, bacteria must counter the changes to their redox state and maintain a strong reducing intracellular environment. Bacteria employ two primary strategies to achieve redox homeostasis. The first involves direct enzymatic detoxification of small reactive molecules, such as through the conversion of superoxide to hydrogen peroxide by superoxide dismutase, or the breakdown of hydrogen peroxide to water by catalase (5). The second strategy involves maintaining millimolar concentrations of low molecular weight (LMW) thiols such as glutathione (GSH), mycothiol (MSH) or bacillithiol (BSH). At high concentrations typically found in cells, these LMW thiols readily undergo oxidation when exposed to oxidative stressors, thereby buffering the intracellular redox environment and protecting macromolecules from oxidative damage (7). LMW thiols also facilitate the repair of protein disulfides through thiol-disulfide exchange reactions and play an important role in maintaining intracellular metal homeostasis (7, 8). Most bacteria also maintain L-cysteine (L-Cys) as a LMW thiol, but its cytosolic concentration is tightly regulated and maintained low for reasons that are not completely understood.

Previous studies have identified several transcriptional regulators in bacteria that respond to oxidative stress and activate antioxidant responses (3, 9). Among these transcription factors, the global regulator Spx is conserved in low G+C gram positive organisms and regulates the expression of several genes involved in thiol homeostasis (3, 10). Spx is also unique among transcriptional regulators as it has a cryptic DNA binding region that becomes exposed only after conformational changes following oxidation of its thiol-based redox switch and interaction with the C-terminal domain of the RNA polymerase α subunit (11). The latter interaction forms a Spx-dependent transcriptional activation complex (Spx-TAC), which recognizes and engages conserved DNA motifs at the −44 promoter element (11, 12). In addition to Spx activation through oxidation of its redox switch, the intracellular abundance of Spx is also a critical determinant of its function (13). Spx is constitutively transcribed, and its protein levels are controlled by ClpXP-dependent proteolysis (5, 14). This turnover is mediated by the adaptor protein YjbH, which facilitates the recognition and delivery of Spx to ClpXP (14). Under oxidative stress conditions, YjbH undergoes aggregation and thereby prevents Spx degradation, leading to its accumulation within cells (15, 16).

Here, we focus on the Spx homolog in *S. aureus*. While its role as a general stress regulator in *S. aureus* has been recognized (17), the functional significance of signaling through its redox switch is much less appreciated. We previously showed that Spx-dependent response to β-lactam-induced cell wall stress occurs independent of its redox switch. In this study, we define the role of the Spx redox switch in modulating Spx function during disulfide stress. We show that the redox switch is critical for resistance to disulfide stress and neutrophil killing. While inactivation of the redox switch had minimal impact on transcription in the absence of stress, diamide-induced disulfide stress altered the transcription of a quarter of the *S. aureus* protein-coding genes. Unexpectedly, our findings demonstrate that resistance to disulfide stress depends more on tightly regulating L-CySS uptake and maintaining intracellular Fe(II) levels than on other canonical mechanisms attributed to enhancing thiol homeostasis.

## Results

### The Spx redox switch contributes to disulfide stress resistance

The Spx ortholog in *S. aureus* is 80% identical to *B. subtilis* Spx and contains all three of its key characteristic identifiers **(Figure 1A)**, including the CXXC redox switch, the Gly52 residue important for interacting with the α-CTD of the RNA polymerase, and the RPI motif which contributes to DNA binding, Cys10 reactivity, and structural stability (10).

**Figure 1:**
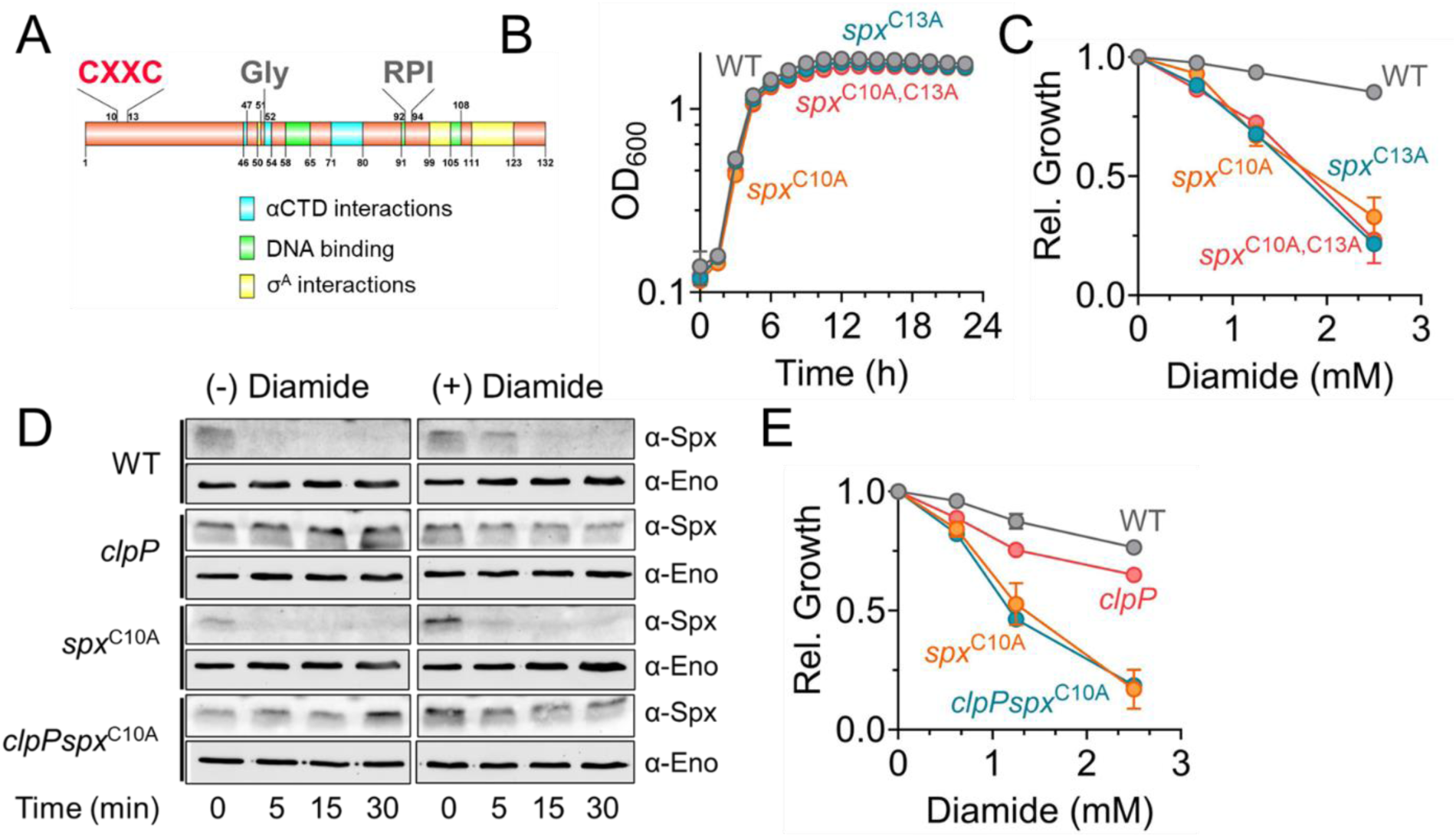
Redox activation of Spx is critical for diamide-induced disulfide stress resistance. **A.** Characteristic features of the *S. aureus* Spx protein **B.** Growth curve of *S. aureus* (WT) and *spx* redox switch mutants. **C.** and **E.** Dose-dependent impact of diamide on growth of *S. aureus*. The relative growth (y-axis) represents the ratio of the area under the growth curve (AUC) for cultures treated with increasing diamide concentrations to that of the untreated control ((−) diamide). **D.** Spx turnover was assessed by western blot analysis using anti-Spx antibodies. Protein translation was inhibited by 100 µg/mL chloramphenicol prior to determining protein turnover. For all experiments, n= 3. Data represents, mean ± SD. Error bars for some data points may not be visible due to small error values.

A previous report suggested that the *spx* allele is essential in *S. aureus*, as its deletion was only possible in the presence of a *rpoB* suppressor mutation or when either thioredoxin or thioredoxin reductase was expressed *in trans* (18). Consistent with this report, our attempts to create an in-frame *spx* deletion in *S. aureus* JE2 were also unsuccessful. Unexpectedly, we were able to introduce point mutations in the Spx redox switch (Cys-X-X-Cys, **Figure 1A)** with relative ease, replacing Cys10, Cys13, or both with Ala (*spx*^C10A^, *spx*^C13A^, and *spx*^C10,13A^) demonstrating that the redox switch of Spx is dispensable in *S. aureus*. Furthermore, the redox switch mutants did not exhibit any growth defects compared to the wildtype (WT) strain **(Figure 1B)**.

Since the Spx redox switch senses and responds to thiol-specific oxidative stress, we challenged the WT, *spx*^C10A^, *spx*^C13A^, and *spx*^C10,13A^ mutant strains with diamide, a well characterized electrophilic stressor that oxidizes cellular thiols. Compared to the WT strain, mutants expressing all three Cys→ Ala Spx variants were hypersensitive to diamide in a dose-dependent manner **(Figure 1C)**, indicating that the Spx redox switch is critical for countering diamide-induced thiol oxidation. Since the diamide sensitivity of all three mutants were similar, we performed all subsequent experiments using the *spx*^C10A^ mutant strain alone. Complementation of the *spx*^C10A^ mutant with the wildtype *spx* allele in the SaPI locus restored diamide resistance **(Figure 1-figure supplement 1)**, suggesting that the observed sensitivity to diamide was not due to off-target effects in the mutant.

### Inactivation of the Spx redox switch increases its turnover but is not the cause of disulfide stress sensitivity

In *S. aureus*, Spx is constitutively transcribed, but its levels are kept low as it is continuously degraded by the intracellular ClpXP protease (5). However, Spx stability is predicted to increase under various stress conditions, leading to transcriptional activation (15). To determine if the oxidation of the Spx redox switch impacts its stability, we measured the turnover of Spx and the Spx^C10A^ variants following exposure of the WT and *spx*^C10A^ mutant to 1 mM diamide, respectively **(Figure 1D)**. Protein synthesis in the WT and *spx*^C10A^ mutant was arrested by chloramphenicol treatment immediately prior to assessing Spx turnover. Both the native and the Spx^C10A^ variant exhibited rapid degradation, with half-lives of 1 min and 0.68 min, respectively **(Figure 1D, Figure 1-figure supplement 2)**. Diamide treatment of the WT strain led to an ∼8.6-fold increase in Spx stability (half-life 8.69 mins) **(Figure 1D, Figure 1-figure supplement 2)**. However, the stability of the Spx^C10A^ variant under diamide stress only increased by ∼5-fold **(Figure 1D, Figure 1-figure supplement 2)**. These results suggest that differences in Spx stability between the WT and *spx*^C10A^ mutant following diamide challenge may account for the latter mutant’s increased diamide sensitivity.

To test if decreased stability of Spx^C10A^ protein variant contributed to the diamide sensitivity, we inactivated *clpP* in the *spx*^C10A^ mutant background. We hypothesized that loss of *clpP* function would prevent Spx^C10A^ degradation leading to increased diamide resistance. As expected, *clpP* inactivation resulted in the stabilization of both Spx and Spx^C10A^ in both the WT and *spx*^C10A^ mutant, respectively, for over 30 minutes **(Figure 1D)**. However, despite increased stability of Spx^C10A^, the *clpPspx*^C10A^ double mutant remained sensitive to diamide (compare *spx^C10A^* to *clpPspx^C10A^* double mutant; **Figure 1E)**. These results suggest that the changes in Spx^C10A^ turnover or its intracellular abundance cannot account for the increased diamide sensitivity of the *spx^C10A^* mutant. Instead, oxidation of the Spx redox switch itself appears to be crucial for diamide resistance, likely due to direct modulation of genes involved in thiol homeostasis.

### BSH redox response to diamide is similar in the WT and *spx^C10A^* strains

To gain insight into the basis of diamide hypersensitivity in the *spx^C10A^* mutant, we compared transcriptional profiles of the WT and *spx^C10A^* strains before and after diamide challenge **(Figure 2A,B)**. RNA sequencing analysis (RNA-seq) of exponentially grown cultures revealed very few changes to gene expression (28 genes) between the WT and *spx^C10A^* mutant in the absence of diamide **(Figure 2A)**. However, upon diamide challenge, the *spx^C10A^* mutant had 715 genes that were differentially regulated and significant at an adj *P-*value of 0.05 **(Figure 2B)**. Among these, 237 genes were upregulated and 124 were downregulated by at least two-fold. KEGG pathway analysis indicated that multiple metabolic pathways were impacted in the *spx^C10A^*mutant **(Figure 2-figure supplement 1)**. Notably, expression of genes within the sulfur relay system, menaquinone biosynthesis, glycerolipid metabolism, and the pentose phosphate pathway were downregulated, whereas riboflavin biosynthesis and aromatic compound degradation pathways were upregulated following diamide challenge of the *spx^C10A^* mutant. These findings suggest that the inability of *spx^C10A^* to respond to disulfide stress may have broad physiological consequences.

**Figure 2:**
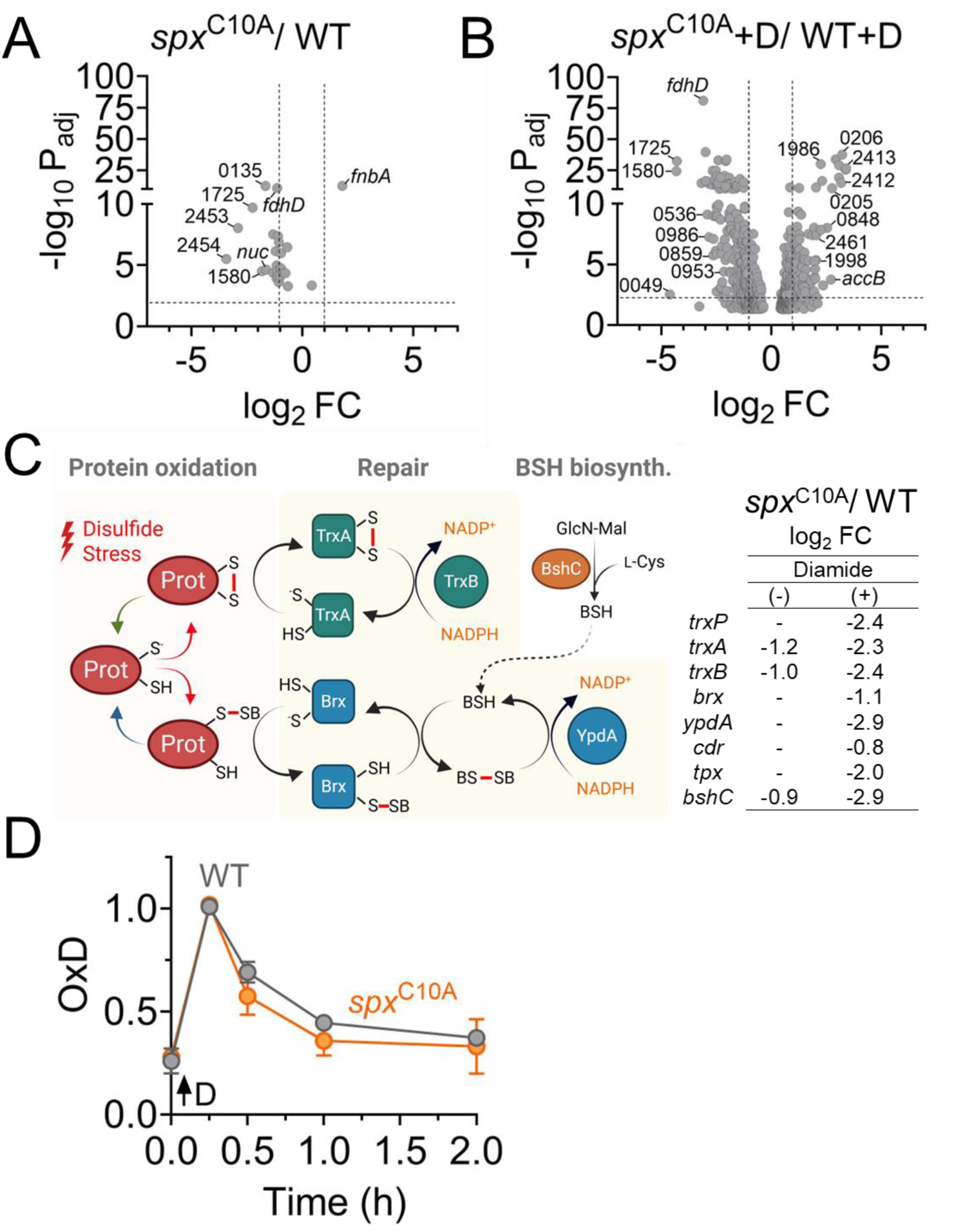
The *spx*^C10A^ mutant efficiently adapts to disulfide stress. **A.** Volcano plot of differentially expressed genes (DEGs) in the *spx*^C10A^ mutant relative to WT strain. **B.** DEGs in the *spx*^C10A^ mutant relative to WT strain after 15 mins of diamide treatment (0.5 mM). +D, diamide treatment. **C.** Expression of thiol-disulfide exchange pathway genes in the *spx*^C10A^ mutant following diamide treatment. **D.** Degree of oxidation (OxD) of bacillithiol over time following 1mM diamide challenge. OxD was assessed using Brx-roGFP2 biosensor as described in material and methods. n= 4, mean ± SD. For all experiments unless otherwise noted, n= 3.

RNA-seq analysis also revealed that several genes from the thioredoxin and bacilliredoxin pathways which participate in protein disulfide reduction and de-bacillithiolation, as well as *bshC*, the enzyme required for BSH biosynthesis, were downregulated in the *spx^C10A^*mutant **(Figure 2C)**. Thus, we hypothesized that diamide stress may alter the redox balance of LMW thiols in the *spx^C10A^* mutant. To test this hypothesis, we used the Brx-roGFP2 biosensor to monitor the redox potential of BSH (*E_BSH_*) over time in both the WT and *spx^C10A^* mutant strains following diamide challenge (1 mM) (19). In the WT strain, diamide rapidly oxidized BSH, shifting *E_BSH_* from –293 mV to –251 mV within 15 minutes **(Figure 2D)**. The oxidation of BSH then gradually reversed and returned to baseline by 2 hours **(Figure 2D)**. Surprisingly, the *spx^C10A^* mutant exhibited a remarkably similar kinetic profile of BSH reduction **(Figure 2D)**, indicating that it is as competent as the WT strain in maintaining thiol homeostasis, despite altered gene expression of its thiol-maintenance pathways **(Figure 2C)**. Thus, these results suggest that the diamide hypersensitivity of the *spx^C10A^* mutant is unlikely to be caused by an imbalance in LMW thiol homeostasis.

### The Spx redox switch helps *S. aureus* bypass L-CySS toxicity under diamide stress

The lack of a difference in *E_BSH_* of the WT and *spx^C10A^* mutant under diamide stress suggested that the mutant may have an alternative strategy to maintain its BSH redox potential. A search for enriched regulons among the differentially regulated genes of the *spx^C10A^* mutant under diamide stress revealed significant upregulation of the CymR regulon **(Figure 3A)**, Since CymR is a master regulator that controls L-CySS/L-Cys uptake and biosynthesis, we hypothesized that enhanced L-CySS uptake by the *spx^C10A^* mutant and its subsequent reduction in the cytoplasm to L-Cys might help buffer changes in the redox status of BSH. Indeed, two specific lines of evidence suggested that the *spx^C10A^* mutant was able to increase L-CySS uptake under diamide stress. First, the total intracellular thiol content significantly increased in the *spx^C10A^* mutant compared to the WT strain under diamide stress **(Figure 3B)**. Importantly, this increase was dependent on L-CySS/L-Cys transporters TcyABC and TcyP (see *tcyA/P* double mutant), suggesting that the *spx^C10A^*mutant imported more L-CySS than the WT following diamide challenge **(Figure 3B)**. Second, the *spx^C10A^* mutant was disproportionately more sensitive than the WT strain to increasing concentrations of selenocysteine, a toxic analog of L-Cys **(Figure 3C)**, whereas inactivation of both L-CySS/L-Cys transporters in the *spx^C10A^*mutant enhanced resistance of *S. aureus* to selenocysteine **(Figure 3C)** Together, these results support increased uptake of L-CySS by *spx^C10A^* mutant under diamide stress. It is important to note here that both the total thiol content measurements and the selenocysteine bioassay indicated that diamide treatment also enhanced L-CySS uptake in the WT strain, although the extent of uptake was lower than the *spx^C10A^* mutant **(Figure 3B, C)**.

**Figure 3:**
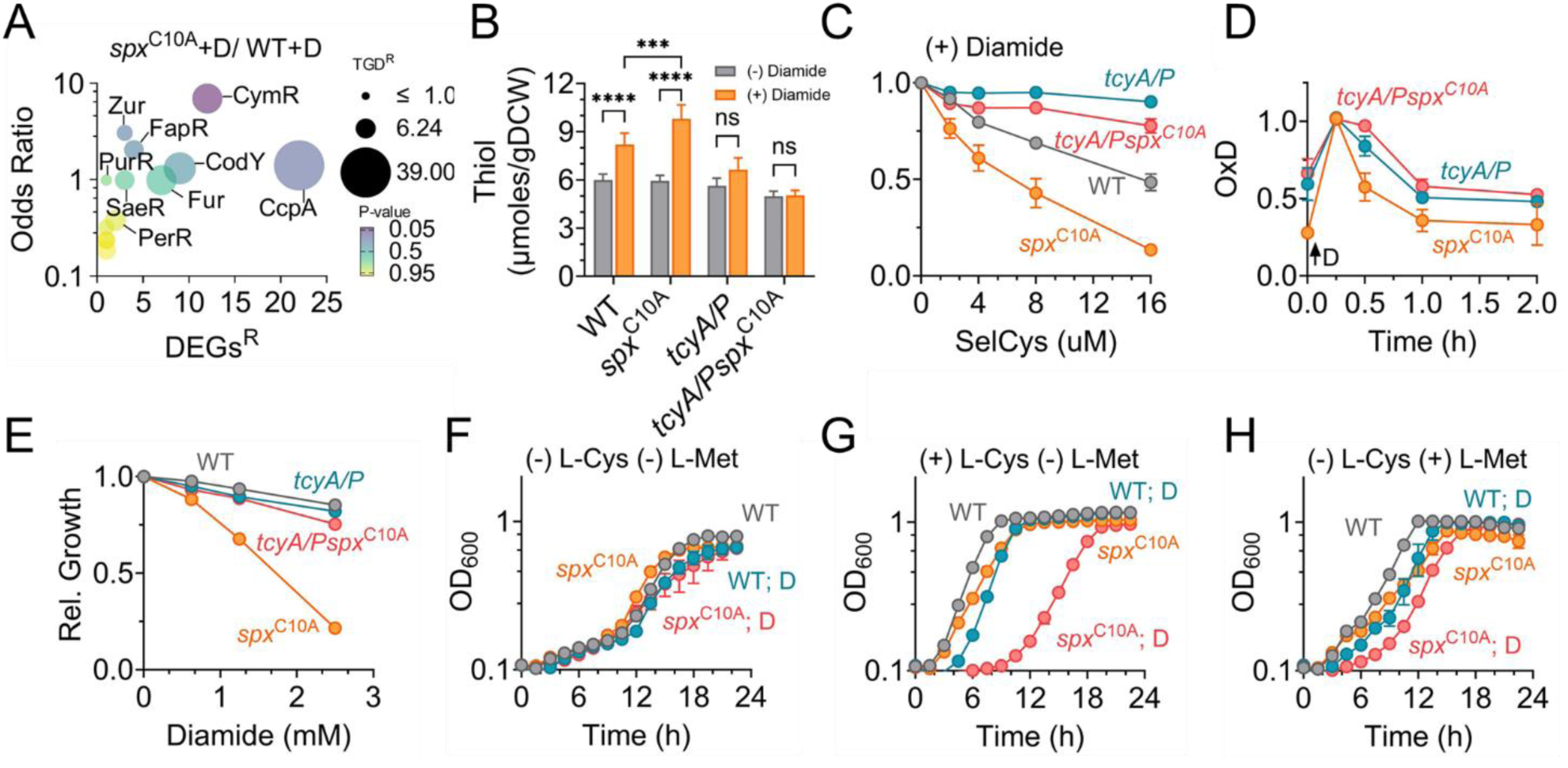
L-Cys accumulation in *S. aureus* reduces disulfide stress but impairs growth. **A.** Bubble plot of enriched regulons in the *spx*^C10A^ mutant relative to WT strain after 15 mins of diamide treatment (0.5 mM). DEGs^R^, total number of differentially expressed genes in the regulon; TGD^R,^ total number of genes designated as part of the regulon; +D, diamide treatment. **B.** Low molecular weight thiol content in cells was determined using the DTNB assay as described in Materials and Methods. gDCW, gram dry cell weight. n= 6, mean ± SD, Two-way ANOVA, Tukey’s post-test. **C.** Dose-dependent impact of selenocysteine on growth of different *S. aureus* strains challenged with 1 mM diamide. **D.** Degree of oxidation (OxD) of bacillithiol as a function of time following diamide treatment (1mM). Diamide (D) was treated at time 0. n= 4. **E.** Dose-dependent impact of diamide on growth of *S. aureus*. **F.**, **G.** and **H.** Growth of *S. aureus* strains in chemically defined media (CDM) with different combinations of sulfur-containing amino acids. For all experiments unless otherwise noted, n= 3. Data represents, mean ± SD, \*\*\**P* ≤ 0.001, \*\*\*\**P* ≤ 0.0001.

Interestingly, inactivating the L-CySS/L-Cys transporters in WT and *spx^C10A^* strains increased BSH oxidation **(Figure 3D**, at time 0), corresponding to a change in *E_BSH_* from −293 mV to −273 mV, indicating L-CySS/L-Cys import as a typical strategy *S. aureus* undertakes to sustain *E_BSH_* at a more reduced state. More importantly, the *tcyA/Pspx^C10A^* mutant, much like the *tcyA/P* mutant, failed to restore *E_BSH_* over time under diamide stress **(Figure 3D)**. Collectively, these results indicate that diamide-induced disulfide stress increased L-CySS uptake in the *spx^C10A^* mutant to re-establish the BSH redox equilibrium.

Remarkably, inactivation of the L-CySS/L-Cys transporters in the *spx^C10A^* mutant (*tcyA/Pspx^C10A^*) rescued its diamide hypersensitivity and conferred resistance **(Figure 3E)**. These results indicate that increased L-CySS uptake under diamide stress is very detrimental to the growth of the *spx^C10A^*mutant despite contributing to the restoration of LMW thiol homeostasis. Notably, the diamide hypersensitivity observed in the *spx^C10A^* mutant specifically resulted from L-CySS uptake (see (+) L-CySS (-) L-Met), as the growth of the WT and *spx^C10A^* mutant was comparable under diamide stress when grown in defined media supplemented with other sulfur sources, such as L-Met (see (-) L-CySS (+) L-Met) or inorganic sulfur salts (see (-) L-CySS (-) L-Met, **Figure 3F-H)**. Together, these findings suggest that increased L-CySS uptake in the *spx^C10A^* mutant forces a trade-off between maintaining thiol balance and supporting growth. But a functional Spx redox switch allows *S. aureus* to avoid this trade-off and maintain thiol homeostasis without excessive L-CySS uptake.

### Toxicity of L-CySS to growth results from decreased bioavailable iron

Next, we examined the nature of the toxicity linked to L-CySS uptake in the *spx^C10A^*mutant under diamide stress. RNA-seq analysis of both the WT and *spx^C10A^*mutant revealed that diamide treatment induced the expression of multiple genes within the Fur regulon **(Figure 4A** and **Figure 4-figure supplement 1).** In addition, regulon enrichment analysis also indicated activation of the Fur regulon in the WT strain, suggesting that iron homeostasis may be affected by diamide-induced disulfide stress **(Figure 4B)**. To test this hypothesis, we measured total intracellular iron levels in both the WT and *spx^C10A^*mutant strains after diamide challenge using ICP-MS **(Figure 4C)**. The *fur* mutant, which is known to accumulate iron, was used as a control for ICP-MS experiments **(Figure 4C)**. Our results showed that while both WT and *spx^C10A^* mutant accumulated iron under diamide stress, the *spx^C10A^* mutant exhibited a greater increase in intracellular iron content **(Figure 4C)**.

**Figure 4:**
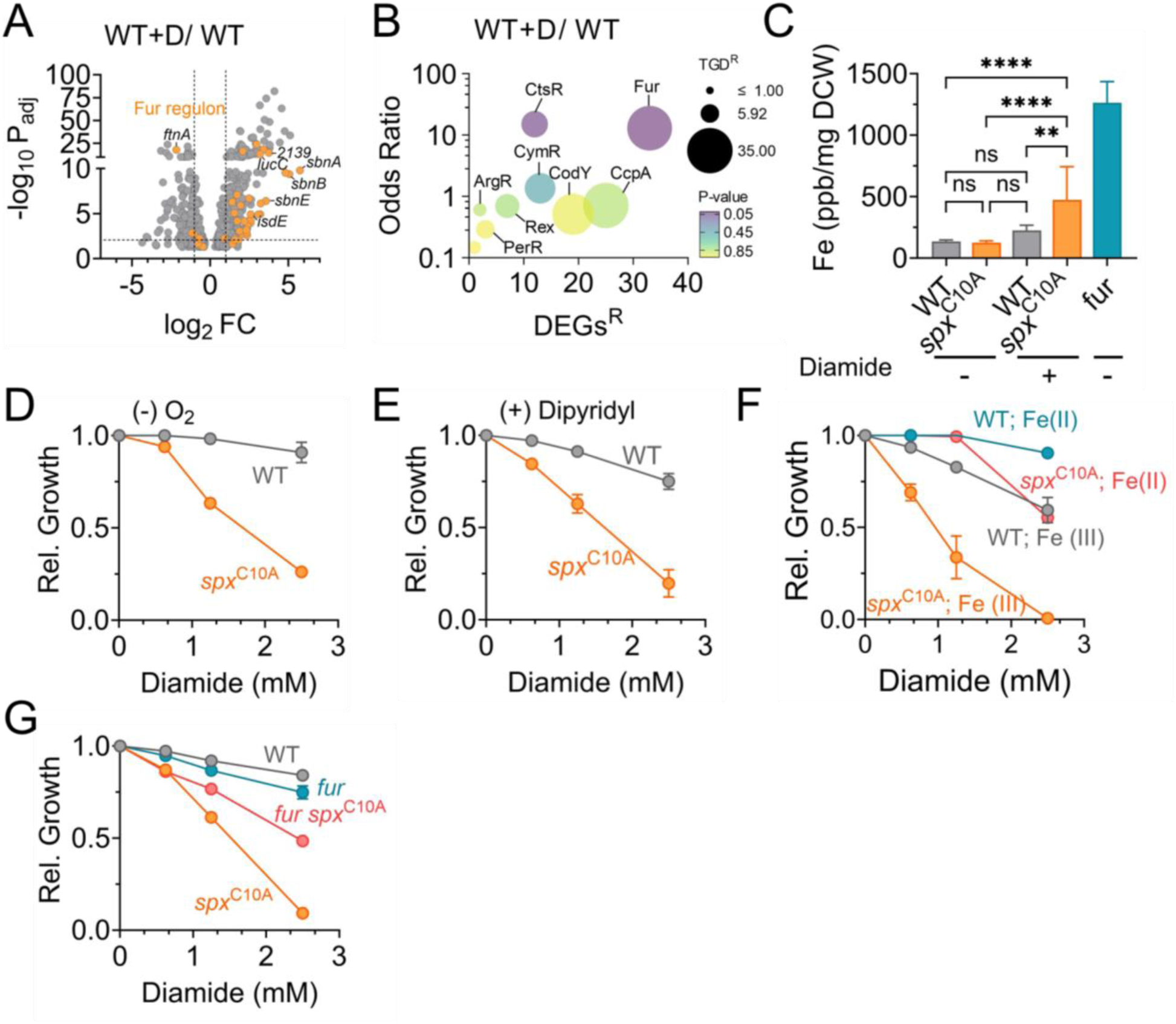
Disulfide stress mediates Fe(II) starvation in *S. aureus*. **A.** Volcano plot of DEGs following diamide-induced disulfide stress in WT strain. +D, diamide treatment. **B.** Bubble plot of enriched regulons in the WT strain following diamide treatment. DEGs^R^, total number of differentially expressed genes in the regulon; TGD^R,^ total number of genes designated as part of the regulon. **C.** ICP-MS analysis of iron pools in *S. aureus*. DCW, dry cell weight. n= 9, mean ± SD, Tukey’s post-test. **C.** Dose-dependent impact of diamide on growth of *S. aureus* strains under **D.** anaerobic **E.** dipyridyl (0.31 mM) **F.** Fe(II) and Fe(III) and **G.** aerobic conditions. For all experiments unless otherwise noted, n= 3, Data represents, mean ± SD. \*\**P* ≤ 0.01, \*\*\*\**P* ≤ 0.0001.

Thus, we initially suspected that the toxicity resulting from increased L-CySS uptake in the *spx^C10A^* mutant under diamide stress may arise from its synergy with elevated intracellular iron, potentially promoting hydroxyl radical generation through Fenton chemistry (20). However, neither growth of the *spx^C10A^*mutant under anaerobic conditions **(Figure 4D)**, nor treatment with the iron chelator dipyridyl **(Figure 4E)**, both expected to disrupt Fenton chemistry, were able to rescue the L-CySS-dependent growth toxicity. On the contrary, supplementation of excess iron completely restored the growth of the *spx^C10A^*mutant. Both Fe(II)NH₄SO₄ and Fe(II)Cl₂ supplementation restored growth of the *spx^C10A^* mutant under diamide stress, whereas Fe(III)Cl₃ did not **(Figure 4F)**. Similarly, increasing intracellular iron through inactivation of *fur* in the *spx^C10A^* mutant also partially restored its diamide hypersensitivity **(Figure 4G)**. Together, these results suggest that increased L-CySS uptake and its reduction to L-Cys may have resulted in a decrease in bioavailable Fe(II) in the *spx^C10A^* mutant.

To confirm whether L-CySS uptake impacted Fe(II) levels, we evaluated the sensitivity of the WT, *spx^C10A^*, *tcyA/P* and *tcyA/Pspx^C10A^* mutants to streptonigrin, an antibiotic whose lethality depends on intracellular Fe(II) levels. If bioavailable Fe(II) levels are indeed low in the *spx^C10A^* mutant, we reasoned that this mutant should display increased resistance to streptonigrin following diamide challenge. Furthermore, if this resistance is driven by increased L-CySS uptake, then inactivation of both the L-CySS/L-Cys transporters in the *spx^C10A^*mutant background should restore streptonigrin sensitivity. Consistent with these predictions, we note that the *spx^C10A^* mutant which was pre-exposed to 5 mM diamide was more resistant to streptonigrin, having a higher specific growth rate than the WT strain grown under the same conditions **(Figure 5A, B)**, whereas in the absence of streptonigrin, both the WT and *spx^C10A^* mutant strains exhibited similar growth rates confirming that observed phenotypes were specific to streptonigrin treatment.

**Figure 5:**
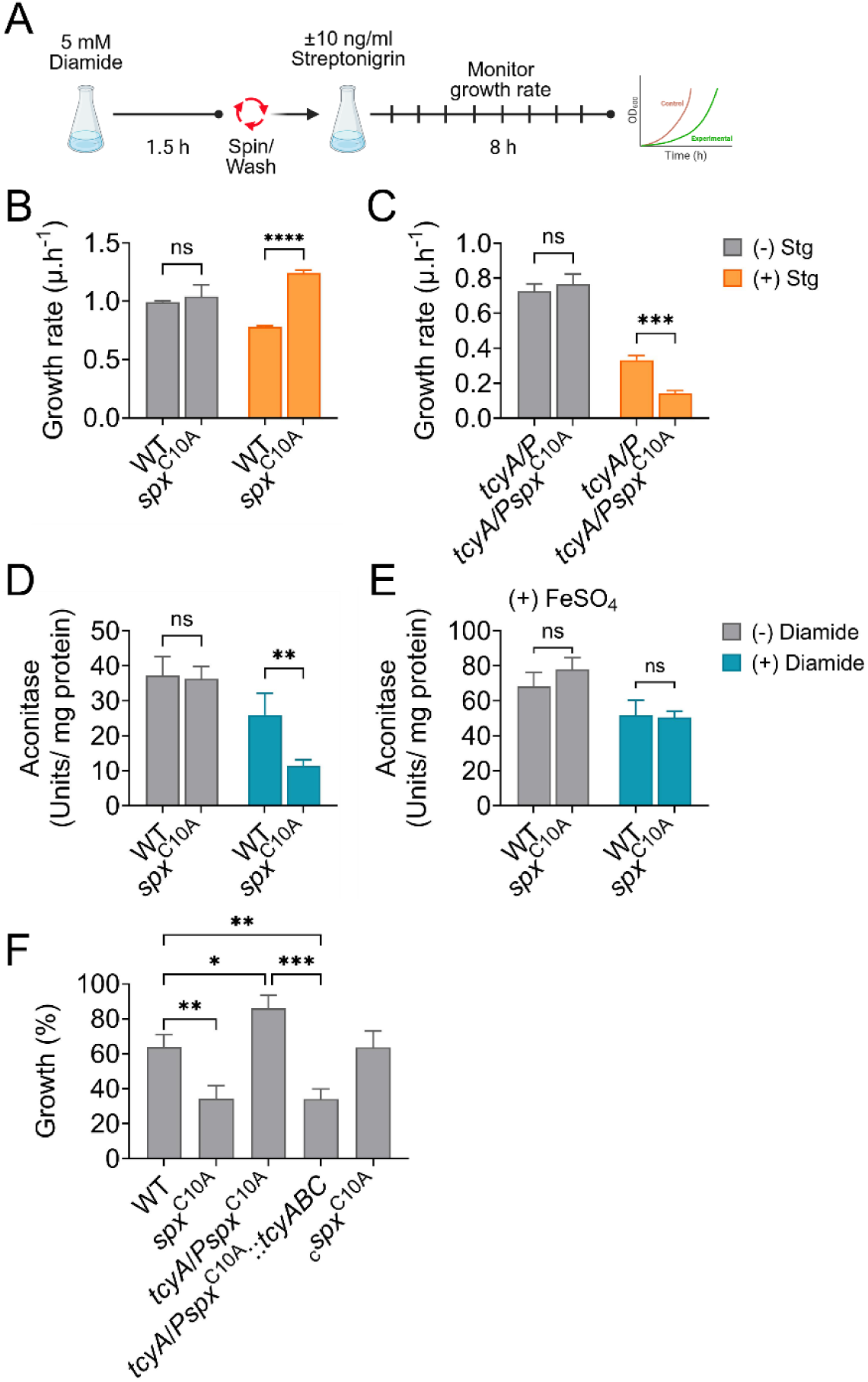
L-Cys uptake in response to disulfide stress mediates Fe(II) starvation. **A.** Schematic depicting timeline of streptonigrin treatment and growth assessment of *S. aureus* cultures after diamide exposure. Diamide was removed prior to streptonigrin treatment. The specific growth rates (µ.h^-1^) of **B.** WT, *spx*^C10A^ and **C.** *tcyA*/*P*, *tcyA*/*Pspx*^C10A^ were determined from growth curves using the equation (*lnN*_2_ − *lnN*_1_)/(*t*_2_ − *t*_1_), where *N*_2_ and *N*_1_ are OD_600_ at times *t*_2_ and *t*_1_. Aconitase activity was measured from either, **D.** crude cell extracts of cultures grown with or without 1 mM diamide or, **E.** from samples grown in media supplemented with 4 mM FeSO_4_. **F.** Neutrophil killing assay. Opsonized *S. aureus* was incubated with neutrophils at an MOI of 1 for 4 h, and bacterial viability was determined. Percent growth in the presence of neutrophils is quantified relative to the strain cultured in the absence of neutrophils (100% = no neutrophil killing, <100% = neutrophil killing, >100% = growth enhancement in the presence of neutrophils). n= 7, mean ± SEM, One-way ANOVA, Tukey’s post-test. For all other experiments, n= 3, mean ± SD, Two-way ANOVA, Tukey’s post-test. \**P* ≤ 0.05, \*\**P* ≤ 0.01, \*\*\**P* ≤ 0.001, \*\*\*\**P* ≤ 0.0001.

In contrast, the inactivation of the L-CySS/L-Cys transporters in the *spx^C10A^* background reversed the streptonigrin resistance phenotype **(Figure 5A, C)**. Indeed, the *tcyA/Pspx^C10A^* mutant was more sensitive to streptonigrin than either the WT or *tcyA/P* mutant, suggesting that the intracellular Fe(II) levels were higher in the *tcyA/Pspx^C10A^* than the other strains tested. Additionally, no difference in the growth rates of the *tcyA/P* and *tcyA/Pspx^C10A^* mutants were observed in the absence of streptonigrin challenge. These findings strongly suggest that the growth defect observed in the *spx^C10A^*mutant following diamide stress is driven by a decrease in bioavailable Fe(II) levels resulting from increased L-CySS uptake.

Low Fe(II) levels could adversely impact the activity of enzymes that rely on Fe(II) as a cofactor. To test this, we measured aconitase activity, a key TCA cycle enzyme whose catalytic function depends on a [4Fe-4S] cluster (21). Because aconitase expression is repressed under glucose-rich conditions due to carbon catabolite repression (22), *S. aureus* was cultured in medium (Tryptic Soy Broth) lacking glucose. The change in growth conditions did not alter the sensitivity of the *spx^C10A^* mutant to diamide, nor did it affect its rescue upon inactivation of L-CySS/L-Cys transporters or supplementation of the medium with Fe(II) **(Figure 5-figure supplement 1)**. Consistent with low intracellular Fe(II) levels under diamide stress, our results show that aconitase activity was significantly reduced in the *spx^C10A^*mutant following diamide challenge **(Figure 5D)**. Moreover, supplementation of Fe(II) in the media restored aconitase activity in the *spx^C10A^* mutant to WT levels **(Figure 5E)**. These findings indicate that increased L-CySS uptake during diamide stress limits bioavailable Fe(II) in the *spx^C10A^* mutant and reduces the activity of Fe-dependent enzymes, which may contribute to its growth defect.

### L-Cys-dependent toxicity reduces the survival of the *spx* mutant in human neutrophils

Host neutrophils generate hypohalous acids that can exert disulfide stress in *S. aureus*. To determine whether the L-Cys-dependent toxicity observed in the *spx^C10A^* mutant under disulfide stress affects its ability to resist neutrophil killing, we incubated opsonized *S. aureus* with human neutrophils from healthy donors and measured bacterial survival. As expected, the *spx^C10A^* mutant was significantly more sensitive to neutrophil killing than the WT strain, a phenotype that was fully complemented by expression of spx *in cis* **(Figure 5F)**. Remarkably, the sensitivity of the *spx^C10A^* mutant was dependent on the uptake of L-CySS **(Figure 5F)**. Whereas inactivation of the L-CySS/L-Cys transporters in the *spx^C10A^*background (*tcyA/Pspx^C10A^*) rescued *spx^C10A^* survival to WT levels, introduction of *tcyABC* back into the *tcyA/Pspx^C10A^*mutant strain *in cis* restored susceptibility to neutrophils to the levels observed in the *spx^C10A^* mutant **(Figure 5F)**. Moreover, the differences in susceptibility to neutrophil killing were not due to variations in neutrophil ROS production, which remained consistent across all strains tested (**Figure 5-figure supplement 2)**. These results strongly suggest that activation of the Spx redox switch and its control of L-CySS uptake are critical for *S. aureus* survival in the presence of human neutrophils.

## Discussion

Disulfide stress is a distinct form of oxidative stress that arises when non-native disulfide bonds form within proteins leading to misfolding and proteotoxic stress (23). In this study, we used diamide, a synthetic thiol-specific oxidant that selectively induces protein disulfide bond formation without broader cytotoxic effects (24), providing a precise means to examine how *S. aureus* senses and adapts to disulfide stress. Broadly, our findings reveal that *S. aureus* adapts to disulfide stress by activating Spx, which plays a critical role in restricting L-CySS uptake into cells.

Although L-CySS uptake can effectively contribute to thiol balance under disulfide stress, we found that its excessive accumulation in the cytosol becomes toxic due to disruption of iron homeostasis. Most importantly, activation of the Spx redox switch and its control over L-CySS uptake are essential for *S. aureus* to withstand neutrophil-mediated killing.

Spx functions as a redox sensor through its conserved CXXC motif (25). When exposed to oxidants such as diamide, a disulfide bond forms between the two conserved cysteine residues (Cys10 and Cys13), allowing it to interact with RNA polymerase and activate transcription from specific promoter regions (10, 12). We recently reported that increasing the abundance of Spx in *S. aureus*, by inactivating the ClpXP protease or the Spx adaptor protein, YjbH, enhanced resistance to β-lactam antibiotics (13). Unexpectedly, β-lactam resistance did not depend on a functional Spx redox switch indicating that not all Spx functions require a redox trigger (13). However, in the current study, we show that the Spx redox switch function promotes *S. aureus* resistance to disulfide stress, whereas Spx abundance is less critical. Thus, *S. aureus* Spx uses distinct mechanisms to protect cells from various stresses, including some that work independent of its redox switch, a characteristic also attributed to other Spx orthologs (26, 27).

How does Spx contribute to disulfide stress resistance in *S. aureus*? The *spx*^C10A^ mutant cannot form an intramolecular disulfide bond required for redox switch activation. Hence, it is unable to induce genes involved in thiol-disulfide exchange and BSH biosynthesis, both important for combating disulfide stress and protecting cells from oxidative damage. This raised the possibility that the intracellular environment of the *spx*^C10A^ mutant might remain more oxidized under disulfide stress thereby contributing to the observed growth delay of this strain upon diamide challenge. However, we did not find any evidence to suggest that the *spx*^C10A^ mutant had a more oxidized intracellular environment compared to the WT strain under disulfide stress. The dynamic changes in the redox potential of the major LMW thiol, BSH, an indicator of the intracellular redox state, was found to be similar in both the WT and *spx*^C10A^ mutant following diamide challenge, indicating that *S. aureus* could rapidly adapt to the loss of Spx activation through compensatory mechanisms that maintain intracellular redox balance. Moreover, the ability of the *spx*^C10A^ mutant to efficiently counter bacillithiol oxidation also indicated that the intracellular environment retained sufficient pools of reduced NADPH to fuel a reducing environment.

Our data now reveals that *S. aureus* compensates for the loss of Spx redox-switch function by importing L-CySS into cells **(Figure 6)**. L-CySS, is rapidly reduced to L-Cys and contributes to the LMW thiol pool once it is imported into the cell (28). In *S. aureus*, the thiol-based regulator CymR, represses L-CySS/L-Cys uptake and biosynthesis during growth (29, 30). Previously, it was reported that oxidation of the conserved Cys25 residue in CymR interferes with its regulatory activity, leading to enhanced L-CySS uptake and accumulation in the cytoplasm (31). Indeed, our transcriptomic and biochemical analyses indicated that the *cymR* regulon is upregulated during disulfide stress and that L-CySS uptake was markedly increased in the *spx*^C10A^ mutant. Inactivation of the L-CySS/L-Cys transporters *tcyA* and *tcyP* in *spx*^C10A^ mutant background eliminated L-Cys buildup in the cytoplasm and prevented recovery of the BSH redox potential, strongly supporting the idea that enhanced L-Cys import compensated for the loss of Spx-dependent thiol homeostasis in the *spx*^C10A^ mutant.

**Figure 6:**
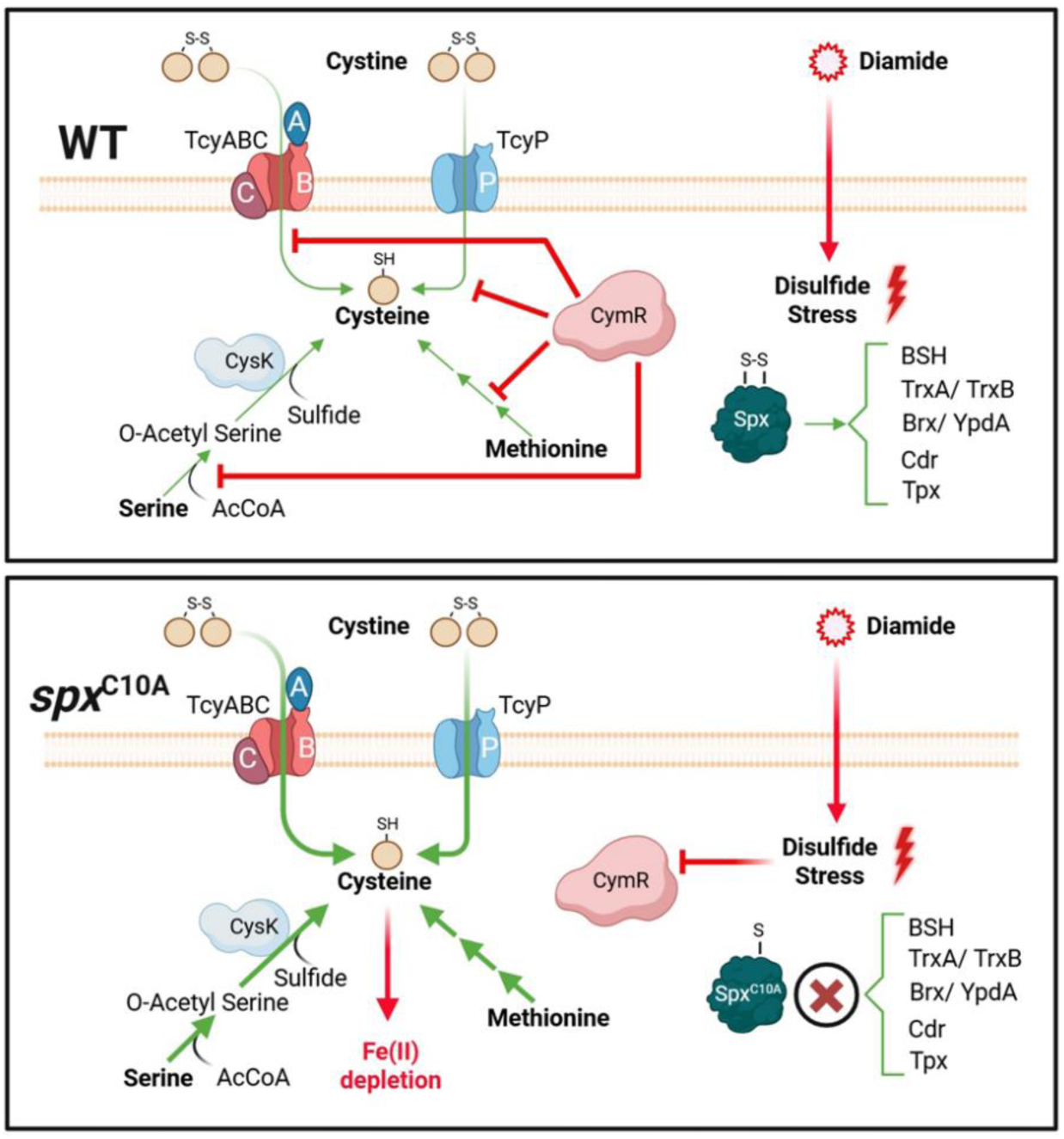
Model depicting regulation of L-Cys uptake by Spx. **Top panel:** L-CySS uptake and biosynthesis are tightly regulated in *S. aureus* by the CymR regulator. Under diamide-induced disulfide stress, Spx is activated through its redox switch and turns on pathways that counter oxidative damage including genes involved in the thiol disulfide exchange system and bacillithiol biosynthesis. A basal level of L-CySS uptake also occurs depending on the level of CymR oxidation which is rapidly converted to L-Cys in the cytosol and contributes to the LMW thiols. **Bottom panel:** Inactivation of the Spx redox switch decreases the ability of *S. aureus* to counter disulfide stress through induction of the Spx regulon. However, under these conditions CymR repression is more strongly relieved due to excess oxidation resulting in increased L-CySS uptake and L-Cys biosynthesis. L-Cys accumulation helps normalize the intracellular redox state. However, excess L-Cys is predicted to chelate Fe(II) resulting in growth inhibition. Thus, *S. aureus* must finely balance Spx activity during disulfide stress to maintain both growth and redox homeostasis.

Surprisingly, we found that disruption of L-CySS uptake through inactivation of *tcyA* and *tcyP* rescued the diamide-induced growth delay of the *spx*^C10A^ mutant. Thus, while increased L-Cys import into the cytoplasm restored redox balance, excessive L-Cys accumulation appeared to reduce *spx*^C10A^ mutant growth during disulfide stress. The toxicity of L-Cys has previously been shown to arise from hydroxyl radical generation via Fenton chemistry when cells are under hydrogen peroxide stress (20). However, our study shows that the mechanism of L-Cys toxicity is fundamentally different when cells are under disulfide stress and involves a decrease in available Fe(II). The basis for Fe(II) limitation under L-Cys-rich conditions is consistent with strong sequestration or chelation of Fe(II) by L-Cys (32–34). In *S. aureus* the cytoplasmic pH is ∼7.8 during aerobic growth and can exceed 8.0 under fermentative conditions (35, 36). The changes to intracellular pH can shift the protonation states of the thiol and α-amine groups of L-Cys and thereby their metal-binding properties (32, 33). With a reported L-Cys-thiol microscopic pKₐ of 8.38 (33), the L-Cys-thiolate fraction is expected to be ∼21% at pH 7.8 and ∼57% at pH 8.5. Thus, an intracellular L-Cys pool close to 0.2 µmol/g dry weight (≈0.13 mM assuming 1.55 mL/g intracellular water (37, 38)), would correspond to ∼27 µM L-Cys-thiolate at pH 7.8 and ∼74 µM at pH 8.5. Given the high stability of the Fe(II)Cys complex (log β₁ ≈ 6.2), L-Cys will likely sequester 96% to 99% of the <10 µM cellular labile Fe(II) pool (39, 40), between pH 7.8-8.5. A subset of Fe(II)Cys complex is expected to adopt a S,N-bidentate coordination (i.e. Fe(II) bound to the thiolate sulfur and the deprotonated α-amine of L-Cys), forming a stronger chelate. More importantly, diamide stress-induced increase in intracellular L-Cys would further shift speciation toward bidentate complex and potentially, Fe(Cys)₂ species (41), amplifying the chelate effect and diminishing the intracellular Fe(II) available to metalloenzymes. LMW thiols such as BSH and GSH are known to buffer labile metal pools in the cytoplasm, including zinc and iron, respectively, and BSH has also been implicated in Fe-S cluster biogenesis (42–44). Thus, an increase in the proportion of L-Cys relative to other LMW thiols could therefore disbalance this buffering network, impairing the ability of these thiols to maintain proper metal homeostasis.

Several of our observations support Fe(II) chelation by L-Cys. Foremost, supplementation of Fe(II) in the growth medium rescued the L-Cys-dependent growth defect of the *spx*^C10A^ mutant, indicating reduced intracellular iron availability. Consistent with low Fe(II), several genes belonging to the Fur regulon were upregulated in the *spx*^C10A^ mutant following diamide challenge. Second, inactivation of the *fur* regulator in the *spx*^C10A^ mutant, which increases intracellular Fe(II) pools, significantly restored growth under disulfide stress. Third, the *spx*^C10A^ mutant exhibited increased resistance to streptonigrin, an antibiotic whose bactericidal activity depends on intracellular Fe(II) levels. Notably, this streptonigrin resistance was dependent on L-CySS uptake suggesting that L-Cys limits Fe(II) availability. Finally, aconitase activity decreased in the *spx*^C10A^ mutant under diamide stress when intracellular L-Cys levels were high, consistent with iron limitation, whereas this reduction was reversed by supplementing Fe(II) in the medium.

*S. aureus* will likely experience disulfide stress in several host environments during colonization and infection. The small anion, thiocyanate (SCN^-^) is found in varying concentrations (µM to mM) in blood, airways, nasal passages, gut, as well as other secretions (45, 46). Halides are also abundant, with chloride levels in plasma and gastric fluid typically between 100 and 150 mM (47). Peroxidases found in immune and epithelial cells, including neutrophils, eosinophils, gastric mucosa, and salivary glands, use hydrogen peroxide to oxidize halides and thiocyanate, generating hypohalous and hypothiocyanous acids, respectively, which readily react with protein thiols to form disulfides and oxidized sulfur species such as sulfenic, sulfinic, and sulfonic acids (45). Remarkably, our findings show that activation of Spx and its tight control of L-CySS uptake is critical for *S. aureus* to survive neutrophil-derived oxidative stress. During infection, neutrophils are a major source of hypochlorous, hypobromous, and hypothiocyanous acids through the action of myeloperoxidase (3, 24, 25). These reactive species damage bacterial cells by inducing disulfide stress, damaging cytochromes and Fe-S clusters, and halogenating amino acids (26, 27).

In conclusion, our findings highlight the dual nature of the *S. aureus* intracellular L-Cys pool during disulfide stress. While L-Cys uptake can compensate for impaired LMW thiol regulation and redox state under disulfide stress, its uncontrolled accumulation disrupts iron homeostasis and cellular metabolism. Thus, the Spx redox switch functions to fine-tune BSH function and thiol-disulfide exchange systems while avoiding the toxic accumulation of L-Cys. This regulatory balance underscores the importance of Spx in coordinating redox and metabolic responses that allow *S. aureus* to survive oxidative and disulfide stress.

## Acknowledgements

We thank Prof. Peter Zuber, Oregon Health & Science University, for generously providing us with anti-Spx antibodies and Prof. Haike Antelmann, Freie University Berlin for providing the plasmid encoding Brx-roGFP2. This work was funded by NIH/NIAID R01AI125588 and P01AI083211 (Metabolomics Core) to V.C.T., and NIGMS 1R35GM154838 to A.J.M. We also acknowledge Pre-Doctoral Fellowship support from the American Heart Association (25PRE1369371) and the University of Nebraska Medical Center awarded to A.G.H. We are grateful to the UNMC Pharmaceutical Formulation, Delivery & Characterization Core and the UNMC Bioinformatics and Systems Biology Core for their professional assistance. The funders had no role in the study design, data collection, interpretation, and decision to submit this work for publication. The authors have no conflict of interest to declare.

## Data Availability

The RNA seq data have been deposited to the GEO database with the identifier GSE306799.

## Materials and Methods

### Bacterial strains, plasmids and growth conditions

The relevant bacterial strains and plasmids used in this study are listed in **Supplemental Table 1**. *S. aureus* strains were grown in tryptic soy broth (TSB; Difco Laboratories) or in chemically defined media. Bacterial growth was carried out at 37°C, starting from an initial OD₆₀₀ of 0.06, either in flasks or in 96-well microtiter plates. When cultures were grown in flasks, a 10:1 flask-to-volume ratio and agitation at 250 rpm was maintained to ensure adequate aeration. Microtiter plate cultures were incubated in a Tecan Infinite M200 spectrophotometer at 37°C and maximum agitation. *Escherichia coli* ElectroTen Blue was used for routine plasmid maintenance and propagation, and cultures were incubated aerobically in Luria–Bertani (LB) media. Antibiotics and supplements were added as required. Ampicillin, 100 µg/mL; Erythromycin, 5 µg/mL; Streptonigrin, 10 ng/mL; Diamide, 0.625-2.5 mM; Selenocysteine, 2-16 µM; Dipyridyl, 0.31 mM; FeSO_4_, 4mM; FeCl_2_, 4mM and FeCl_3_, 4mM.

For bacterial growth assays involving various supplements, cultures were grown across a range of supplement concentrations to assess dose-dependent effects on growth. Growth data were analyzed as relative growth, defined here as the ratio of the area under the growth curve (AUC) for the test condition ((+) supplement) to that of the untreated control ((-) supplement). Relative growth values were then plotted as a function of supplement concentration.

### DNA manipulation and strain construction

The primers used in this study are listed in **Supplemental Table 2**. *S. aureus* JE2 and its isogenic mutants were obtained from the Nebraska Transposon Mutant Library (NTML). All transposon insertions were re-transduced into the JE2 background using ϕ11 bacteriophage to eliminate potential suppressor mutations. Double mutants were generated by first replacing the Erm^R^ cassette of the transposon insertion with a Kan^R^ or Spec^R^ cassette, followed by transduction of the modified transposon into the desired NTML strain. The complementation of mutant strains was achieved by integrating the full-length gene, under the control of its native promoter, into the SaPI1 site of the JE2 chromosome using the pJC1111 integration vector (48). Mutations in the *spx* redox switch (*spx*^C10A^, *spx*^C13A^, and *spx*^C10,13A^) were generated as previously described (13).

### Determination of Spx turnover

For determination of Spx turnover, 25 ml cultures of *S. aureus* strains were treated with 100 mg/mL chloramphenicol for 30 seconds to halt protein synthesis and cultures harvested at various time points. Where indicated, 1 mM diamide was added prior to chloramphenicol treatment. The collected samples were washed thrice with PBS and resuspended in the same buffer. Bacterial cells were lysed by bead homogenization, and total protein concentrations were determined using the Bradford assay.

For Western blot analysis, 10 µg of total protein was resolved on a 15% SDS–PAGE gel and transferred onto a nitrocellulose membrane using the semi-dry Pierce Power Blot apparatus (Thermo Fisher). To detect Spx, cross-reactive polyclonal antibodies raised against *Bacillus subtilis* Spx were diluted 1:1,500 in PBS containing 0.01% Tween-20 and 5% skim milk. Goat anti-rabbit secondary antibodies (Invitrogen) were diluted 1:40,000 in the same buffer. Signal detection was performed using the SuperSignal West Femto chemiluminescence kit (Thermo Scientific), and images were captured with an iBright CL1000 imaging system (Invitrogen). Enolase was used as a loading control and detected with anti-enolase polyclonal antibodies (Thermo Scientific) at a dilution of 1:2,500. Spx turnover rates were determined by densitometric analysis.

### RNA extraction, library preparation and RNA sequencing

Bacterial cultures were grown to an OD_600_ of 0.7 and challenged with 0.5 mM diamide (1,1′-Azobis(N,N-dimethylformamide)) for 10 minutes. Cells were quickly cooled on an EtOH/dry ice bath and frozen at −80 °C until extraction of RNA. RNA was isolated from three biological replicates grown on different days: cells were lysed mechanically using the FastPrep machine (MP Biomedicals) and RNA was isolated by the RNeasy mini kit (Quiagen, Valencia, Calif) according to the manufacturer’s instructions. RNA libraries for RNA-seq were prepared as follows: All samples were rRNA depleted using the Ribo-zero Magnetic kit (Illumina Inc.), and residual DNA from RNA extraction was removed using the DNase MAX kit (MoBio Laboratories Inc.). The samples were purified using the standard protocol for CleanPCR SPRI beads (CleanNA, NL) and further prepared for sequencing using the NEBNext Ultra II Directional RNA library preparation kit (New England Biolabs). Library concentrations were measured using Qubit HS DNA assay and library size estimated using TapeStation with D1000 ScreenTape. The samples were pooled in equimolar concentrations and sequenced (2 x 150 bp) on a HiSEQ X platform (Illumina, USA). All kits were used as per the manufacturer’s instructions.

### DTNB Assay

Total cellular LMW thiols were measured as described previously (24). *S. aureus* strains were grown to a OD_600_ of 1.0 and treated with 1 mM diamide for 30 minutes. 25 OD_600_ units of diamide-treated culture sample was collected and brought to a final volume of 30 ml using PBS and centrifuged for 5 minutes at 10,000 × g and the pellet was resuspended in 600 μl sample buffer containing 50% acetonitrile in 20 mM Tris-HCl, pH 8.0 (v/v). The suspension was incubated for 10 min at 60°C, and 500 μl of supernatant was mixed with 500 uL sample buffer and 5 μl of 100 mM 5,5′-dithiobis-(2-nitrobenzoic acid) (DTNB) in dimethyl sulfoxide. The sample was incubated at room temperature for 10 minutes. The A_412_ was measured, and the LMW thiol content was calculated using the molar absorption coefficient, 14,150 M^-1^ cm^-1^ (49).

### Intracellular degree of oxidation (OxD)

The OxD was determined as previously reported (19). Briefly, *S. aureus* strains expressing the Brx-roGFP2 biosensor were grown aerobically to an OD_600_ of 1.0. Fluorescence measurements were taken at 0, 15, 30, 60, and 120 minutes after the addition of 1 mM diamide as previously described. Fully reduced and oxidized controls were prepared with 10 mM DTT and 5 mM diamide, respectively. Brx-roGFP2 biosensor fluorescence emission was measured at 510 nm after excitation at 395 and 475 nm using the Tecan Infinite M200 spectrophotometer. The OxD of the Brx-roGFP2 biosensor was determined for each sample and normalized to fully reduced and oxidized controls (eq. 1). Based on the OxD and *E*^0′^*roGFP2* = −280 mV, the BSH redox potential (E_BSH_) was calculated according to the Nernst equation (eq. 2). Any samples which had a calculated OxD >1, the values were capped at 0.99.

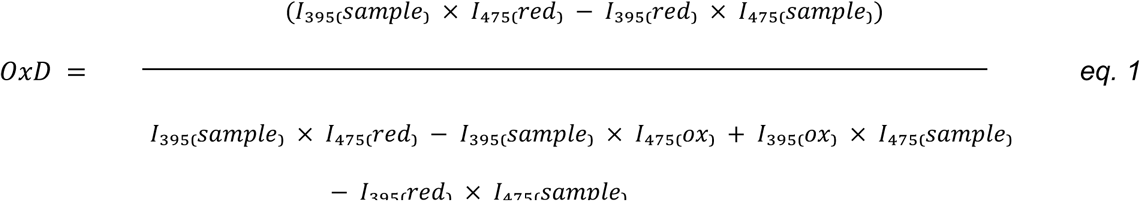

where ‘I’ represents the observed fluorescence excitation intensities.

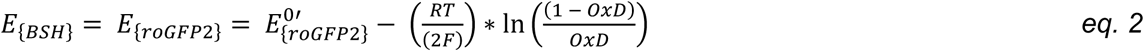

### Inductively coupled plasma mass spectrometry (ICP-MS)

*S. aureus* strains were grown to an OD_600_ of 0.1, then treated with 1 mM diamide for 1.5 hours. Cultures were then harvested and washed by resuspension and centrifugation at 10,000 × g for 10 min, twice with 100 mM EDTA and then twice with milliQ water (18.2 MΩ.cm). Bacterial pellets were desiccated at 96°C overnight. The dry weight of cells was measured, and the pellets were resuspended in 500 uL 35% HNO_3_ (Optima grade) and heated at 95°C for 1 h prior to removal of debris by centrifugation. Samples were diluted 1:10 into a matrix (composed of 2% HNO_3_ and 5 ppb rhodium (internal standard)) and analyzed on a NexION 300Q (Perkin Elmer) spectrometer.

### Aconitase Assay

*S. aureus* was grown aerobically at 37°C in 25 mL TSB without glucose. Cells were harvested and lysed by bead homogenization in lysis buffer containing 90 mM Tris HCl (pH 8.0) and 100 uM fluorocitrate to obtain crude protein lysates. Aconitase reaction was initiated in reaction buffer containing 90 mM Tris HCl (pH 8.0),100 uM fluorocitrate and 20 mM isocitrate by the addition of approximately 100-200 µg of protein from cell lysate. Aconitase activity was measured by monitoring conversion of iso-citrate to cis-aconitate at A_240_ as described previously (50). One unit of aconitase activity is defined as the amount of enzyme necessary to give a ΔA_240_ min^−1^ of 0.0033.

### Streptonigrin sensitivity

*S. aureus* strains were grown to an OD_600_ of 0.2 and treated with 5 mM diamide for 1.5 hours. Cells were harvested by centrifugation, washed once with PBS, and resuspended in TSB to an OD_600_ of 0.06. The cultures were then challenged with streptonigrin (final concentration, 10 ng/mL), and growth was monitored over 24 hours by measuring OD_600_.

### Neutrophil killing assay

Human neutrophils isolated from healthy donors were cultured on a 96-well plate at 2.5 x 10^5^ cells/well in DMEM supplemented with 10% FBS (heat inactivated) and allowed to rest in a 37°C, 5% CO_2_ incubator for 1 hour. Each strain was opsonized in non-heat inactivated FBS at a 1:1 ratio of DMEM:FBS prior to adding to the neutrophils (MOI=1). Cultures went for 4 hours at 37°C in a 5% CO_2_ incubator before cultures were dilution spot plated onto TSA. Colony forming units (CFU) were counted the next morning. Percent *S. aureus* survival was determined by dividing average CFU count from neutrophil containing wells by control wells containing only bacteria (controls).

### Regulon enrichment analysis

Genes corresponding to various *S. aureus* regulons were obtained from the RegPrecise database (51). Only those genes that were present in both the RNA-seq dataset and the RegPrecise regulon database were included in the enrichment analysis. For each regulon, a 2 × 2 contingency table was constructed comparing the numbers of differentially expressed genes (DEGs) and non-DEGs that were either assigned or not assigned to a specific regulon. Enrichment was assessed using the one-sided Fisher’s exact test, implemented in the scipy.stats.fisher_exact function from the SciPy library in Python (52). The odds ratio and P-value were calculated for each regulon.

---

**Figure 1-figure supplement 1:**
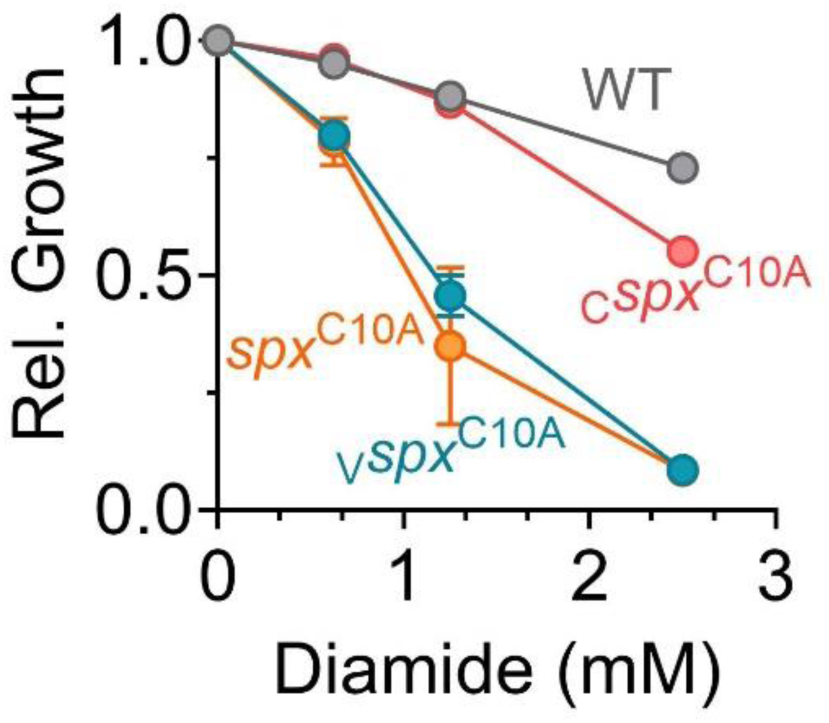
Complementation of *spx*^C10A^. Dose-dependent impact of diamide on growth of various *S. aureus* strains. ***C****spx*^C10A^, *spx*^C10A^ mutant complemented with pJC1111: *spx* at SaPI site, ***V****spx*^C10A^, *spx*^C10A^ mutant with pJC1111 (vector control) at SaPI site; n= 3, Data represent mean ± SD.

**Figure 1-figure supplement 2:**
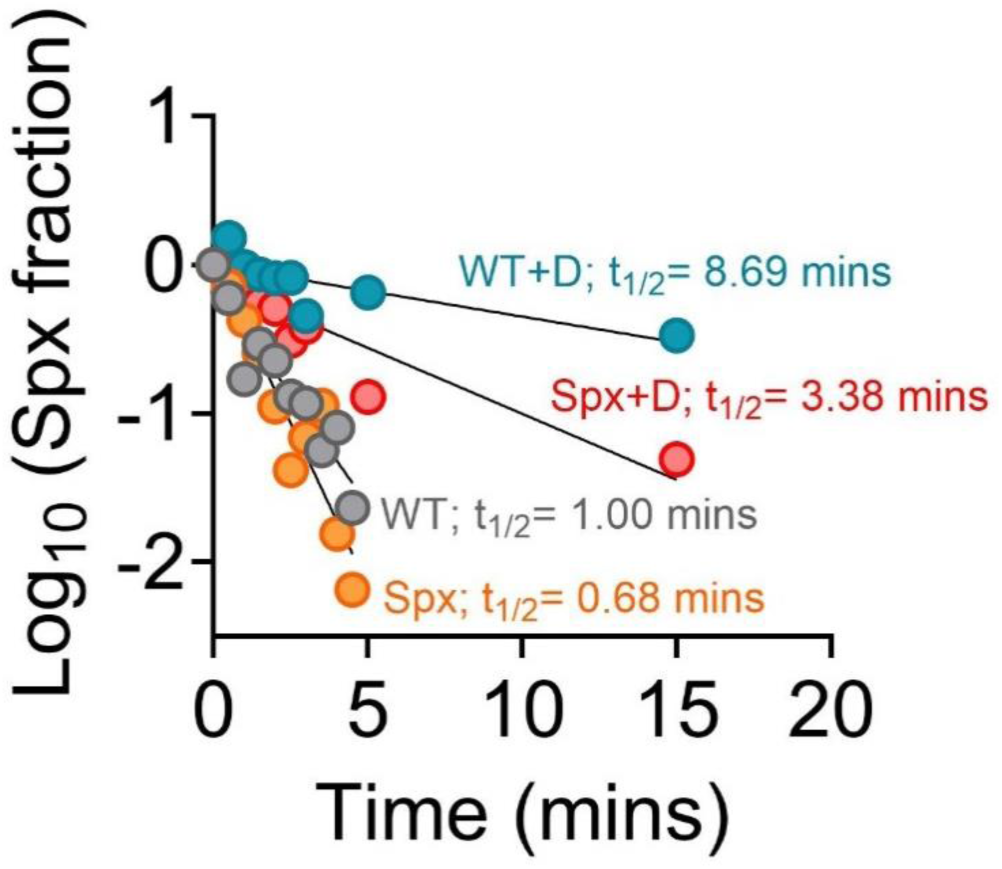
Half-life of Spx. The half-life of different Spx variants were determined from Spx decay intensities observed over 0-15 min for WT+D and *spx*^C10A^+D, and 0-4.5 min for WT and *spx*^C10A^. The Spx intensities derived from western blot analysis was normalized to the loading control (enolase). The half-life (t_1/2_) was calculated as the ratio of Log_10_(0.5) to the slope of the log-transformed protein decay intensity. +D, diamide.

**Figure 2-figure supplement 1:**
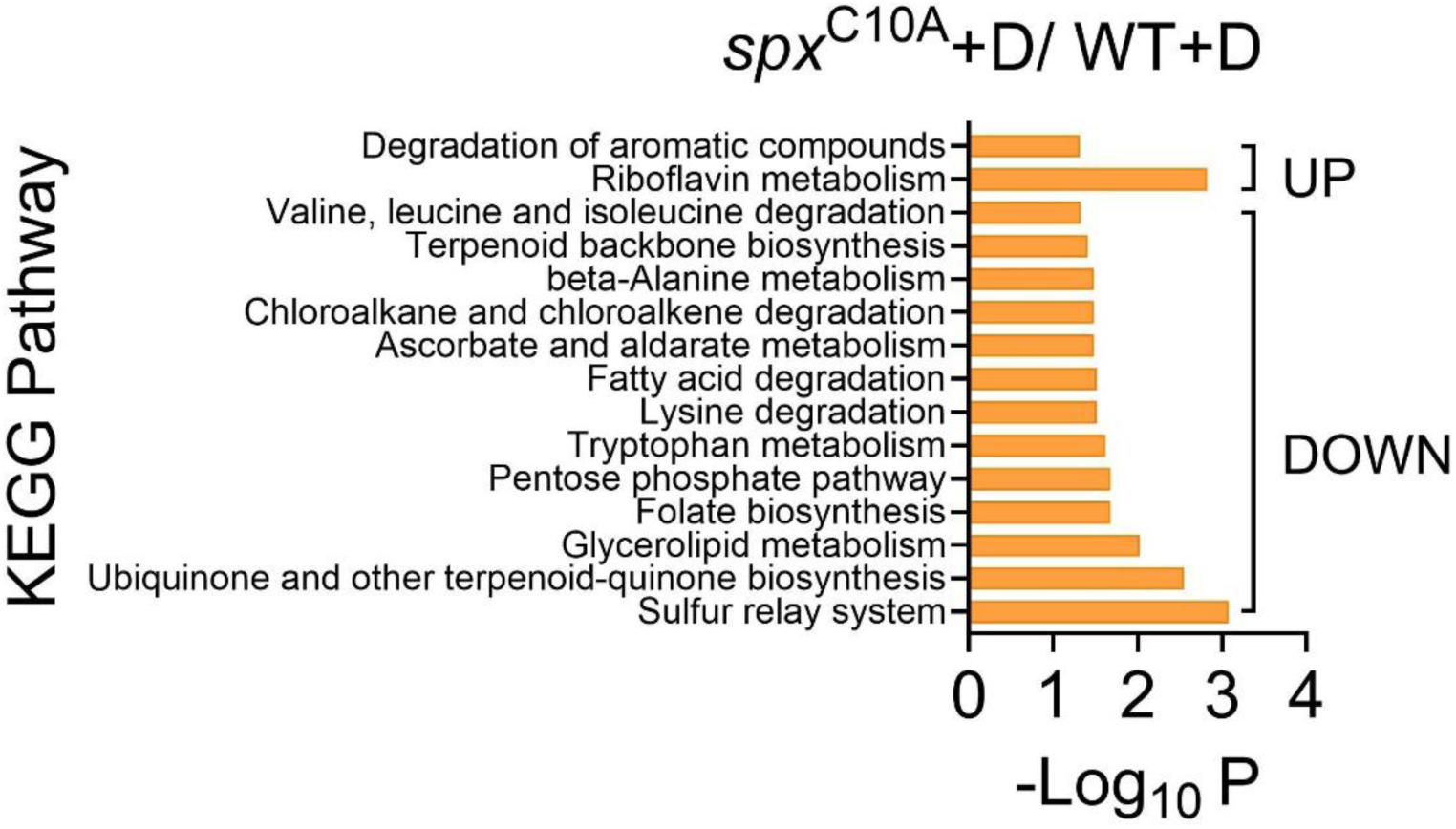
KEGG pathway analysis of RNAseq dataset. Differentially expressed pathways in the *spx*^C10A^ mutant relative to the WT strain after 15 mins of 0.5 mM diamide treatment. Analysis was carried out using KOBAS 3.0.

**Figure 4-figure supplement 1:**
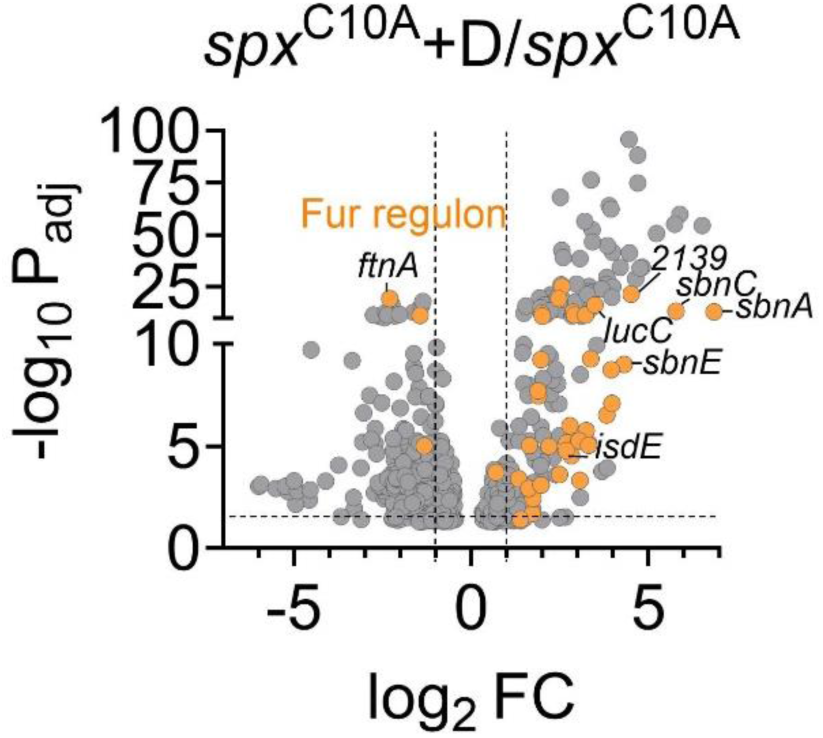
Volcano plot. Differentially expressed genes in *spx*^C10A^ mutant strain following diamide-induced disulfide stress. +D, diamide treatment.

**Figure 5-figure supplement 1:**
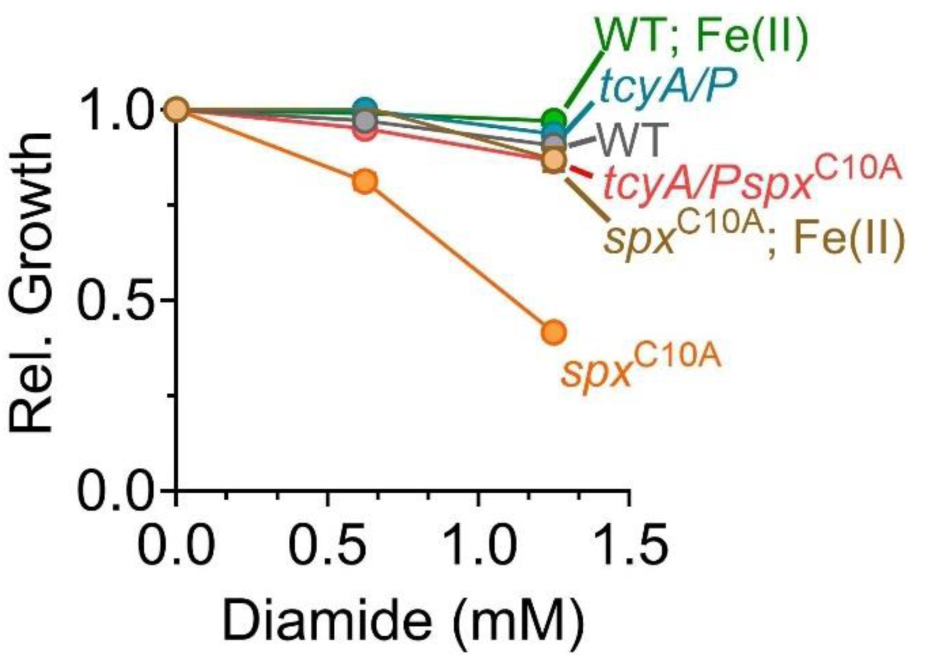
L-Cys-mediated Fe(II) starvation in the *spx*^C10A^ mutant under disulfide stress occurs regardless of glucose in the medium. Cultures were grown in TSB (-) glucose with increasing concentrations of diamide. The relative growth was determined from the fractional area under the growth curve of diamide treated cultures to that of untreated culture.

**Figure 5-figure supplement 2:**
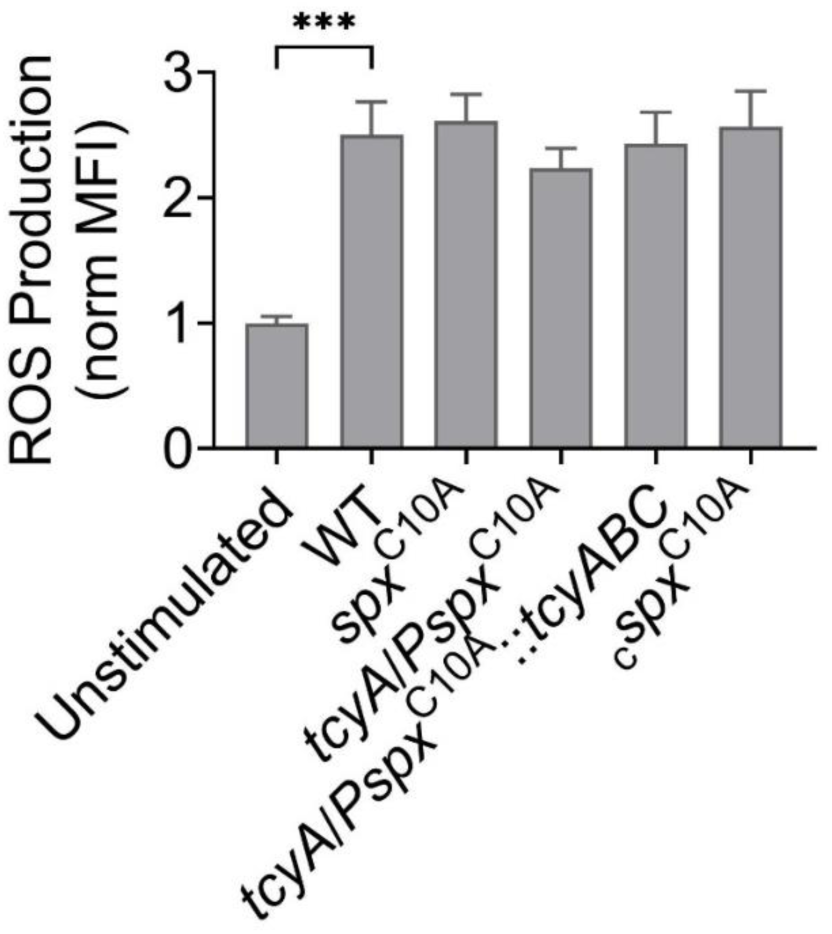
ROS levels in neutrophils. Neutrophils were cultured with *S. aureus* (MOI = 10) for two hours. Neutrophils were stained with DHR123, approximately 15 minutes prior to fixation in 4% PFA. Neutrophils were blocked and stained with anti-CD15 and anti-CD16 antibodies and the mean fluorescence intensity (MFI) for DHR123 in the neutrophil population was quantified by flow cytometry. MFI was normalized to unstimulated neutrophils. n= 5, mean ± SEM, One-way ANOVA, Tukey’s post-test, \*\*\**P* ≤ 0.001.

**Supplemental Table 1:**
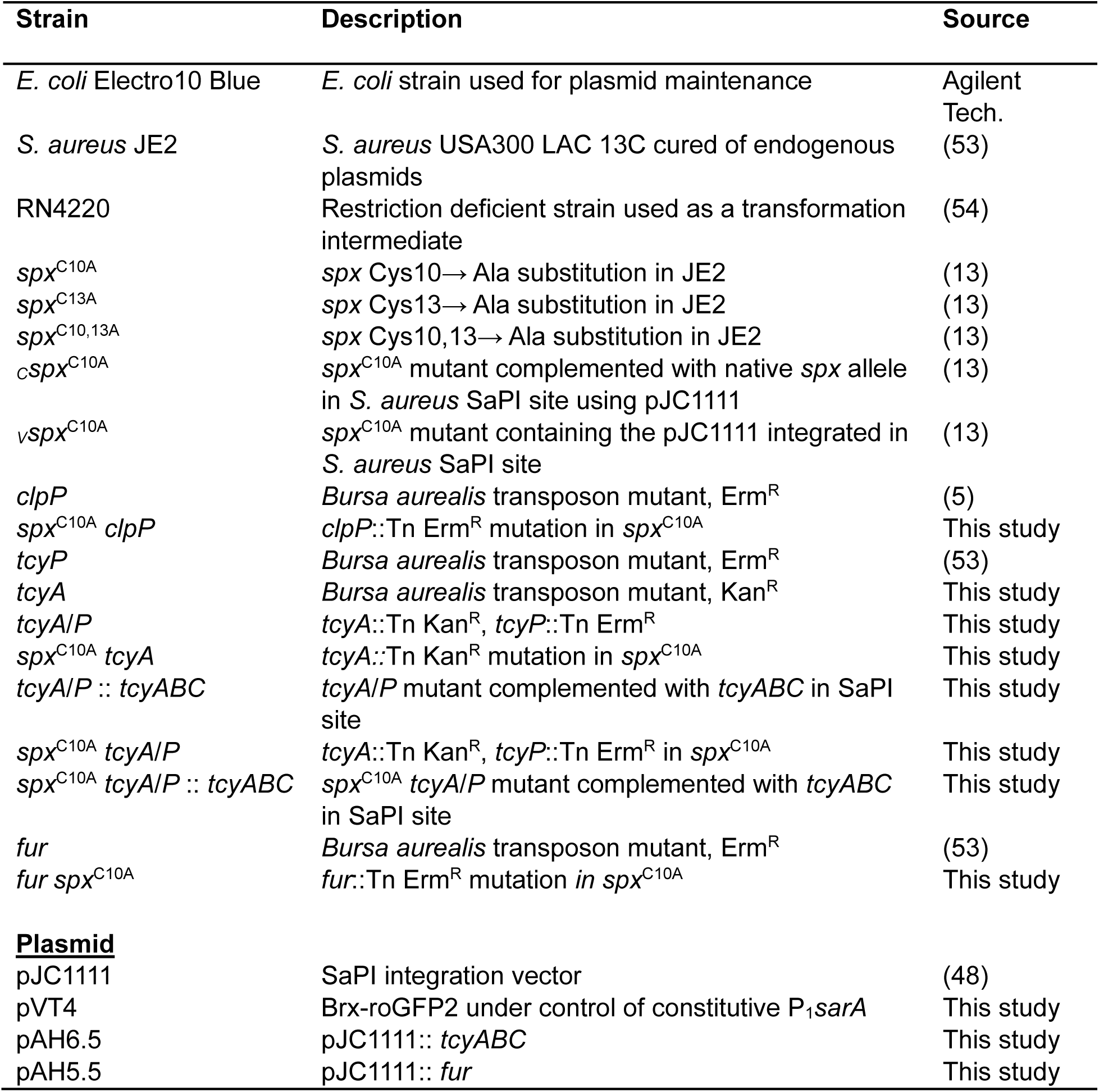
Strains and plasmids used in this study.

**Supplemental Table 2:**
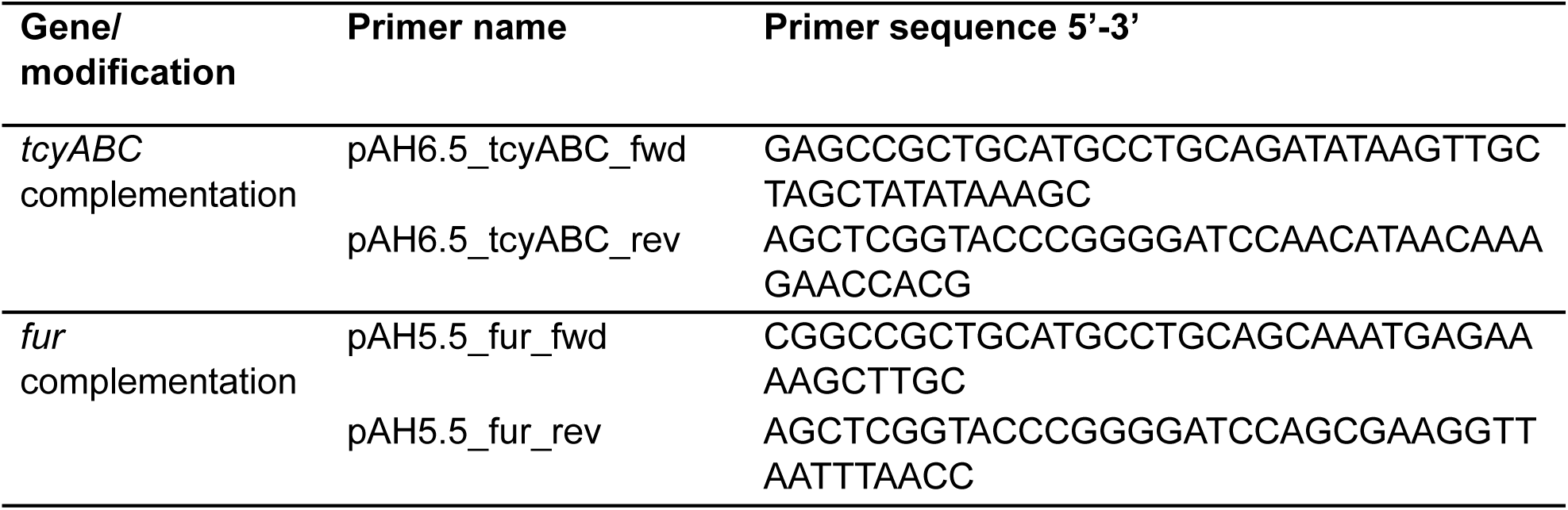
Primers used in this study.

